# Young gene HP6/Umbrea is dispensable for viability and fertility

**DOI:** 10.1101/2023.05.24.542211

**Authors:** Sherilyn Grill, Ashley Riley, Monica Selvaraj, Ruth Lehmann

## Abstract

Studies of the young gene Heterochromatin Protein 6 (HP6) have challenged the dogma that essential functions are only seen in genes with a long evolutionary history. Based on its prominent expression in Drosophila germ cells, we asked if HP6 might play a role in germline development. Surprisingly, we found that CRISPR-generated HP6 null mutants are viable and fertile. We identified an independent lethal allele and an RNAi off-target effect that prevented accurate interpretation of HP6 essentiality in previous studies. We found that the vast majority of young essential genes were viable when tested with orthologous methods. Together our data call into question the frequency with which young genes gain essential functions.

## Introduction

Germ cells have the unique ability to passage an individual’s genome to future generations. This specialized property is essential for the maintenance of most species. In the fruit fly *Drosophila melanogaster*, primordial germ cells (PGCs) are the first cells to form in the developing embryo. From the time of their specification, PGCs must repress somatic differentiation programs to maintain germ cell fate. Despite our understanding of the mechanisms that repress somatic gene expression in PGCs, little is known about how the germ cell-specific transcriptional program is activated or maintained. Previous single-cell RNA-sequencing data from adult, larval, and primordial germ cells have fallen short of identifying a germline transcriptional regulator that is responsible for activating the germline transcriptional program. However, each of these studies detected the putative chromatin regulator *HP6* (also known as *umbrea*) specifically in germ cells (Slaidina et al. 2020; Li et al. 2021; Slaidina et al. 2021). In the germline, *HP6* is zygotically expressed in the PGCs of embryos, the germline stem cells of larval and adult ovaries, and the germ cells of adult testes (Ida et al. 2009; Levine et al. 2012; Slaidina et al. 2020; Li et al. 2021; Slaidina et al. 2021). Despite this, what role HP6 plays in germline development is not known.

HP6 has been characterized as an interactor of the chromatin remodeler HP1 (Greil et al. 2007; Joppich et al. 2009). Its localization to areas of heterochromatin, including the centromeres, telomeres, chromocenter, and the heterochromatic fourth chromosome is dependent on its interaction with HP1 (Greil et al. 2007; Joppich et al. 2009; Ross et al. 2013). In *Drosophila melanogaster*, HP6 has lost the canonical chromodomain of other HP proteins, and instead contains only a chromoshadow domain (Vermaak and Malik 2009; Levine et al. 2012; Ross et al. 2013). The HP6 chromoshadow domain broadly mediates protein-protein interactions, including its interaction with HP1, the HP1 interacting factor HIP, and the telomere capping protein HOAP (Joppich et al. 2009). As HP6 has been shown to localize to areas of heterochromatin, we wondered whether it might play a role in repressing gene expression in the germline to promote germ cell specification.

A potential role for HP6 in germ cell regulation is particularly intriguing because HP6 is a ‘young gene’. Evolutionary studies have suggested that 12-15 million years ago an HP1B gene duplication into an intron of the *dumpy* gene gave rise to the HP6 gene (Vermaak and Malik 2009; Ross et al. 2013). Due to its relatively recent birth, HP6 has been studied as a model for how new genes can acquire essential functions (Chen et al. 2010; Ross et al. 2013). From an evolutionary perspective, essentiality is, in general, associated with deep conservation. How new genes acquire novel functions and whether these functions can quickly become essential is largely unknown (Long et al. 2003; Chen et al. 2010; Chen et al. 2013; McLysaght and Hurst 2016; Kondo et al. 2017). In Drosophila, transgenically-delivered RNAi has been used to interfere with the expression of young genes to identify essential functions (Chen et al. 2010; Xia et al. 2021). However, very few studies have validated these findings with additional genetic analysis. These studies estimated that as many as 30% of young genes acquire essential function, a percentage that has been questioned recently with CRISPR-based mutagenesis studies (Kondo et al. 2017). In only a few cases has the biological function of young essential genes been characterized in detail (Ross et al. 2013; Kasinathan et al. 2020). HP6 is one such example. HP6 was identified as an essential gene in an RNAi screen and was subsequently studied in depth to uncover its centromeric localization in cells (Chen et al. 2010; Ross et al. 2013).

A potential germline specification function for HP6 is particularly interesting in the context of this evolution, as there is significant evidence that new genes can quickly become essential to an organism’s fertility (Loppin et al. 2005; Haerty et al. 2007; Ding et al. 2010; Yeh et al. 2012; Kondo et al. 2017; VanKuren and Long 2018; Witt et al. 2019). Thus, we investigated the biological function of HP6 to determine if HP6 is an essential transcriptional regulator in the germline. Surprisingly, despite previous reports, we show that HP6 is dispensable for viability and fertility. Using genetic analysis, we identified an unlinked lethal mutation in previously generated HP6 mutants and discovered that the observed RNAi lethality is independent of HP6 expression (Chen et al. 2010; Ross et al. 2013). Together, our results demonstrate that HP6 is not essential in *Drosophila melanogaster* and highlight the importance of using multiple orthogonal methods when probing essential gene functions.

## Results

### Generation of an *HP6* null mutants

To test whether HP6 has a function in germline development, we generated *HP6* null mutant alleles by CRISPR gene editing. Two guide RNAs were used simultaneously to target the *HP6* locus near the start codon and stop codon of the HP6 coding region within the first *dumpy* intron. A repair template replaced the *HP6* coding region with the 3xP3-dsRED marker cassette from the pHD-ScarlessDsRed donor vector as previously described (Gratz et al. 2015) (Fig. 1A). dsRED insertion allowed for easy screening of successfully edited animals. Two different founders generated *HP6* deletion alleles (*HP6*^*CR1*^ and *HP6*^*CR2*^), each genetically identical at the *HP6* locus and confirmed by PCR amplification and Sanger sequencing. Isogenic *HP6* mutant lines were generated and kept as balanced stocks. Surprisingly, all HP6^CR^ mutant stocks derived from the original founders contained homozygous mutant flies in addition to heterozygous balanced flies. This suggested that loss of HP6 does not affect viability in *Drosophila melanogaster*. To confirm that *HP6*^*CR*^ null flies contained two copies of the edited HP6 mutant allele, genomic DNA was isolated from individual *HP6*^*CR*^ homozygous mutant or heterozygous animals and the *HP6* locus was amplified by PCR. Amplified products were run on an agarose gel and genotypes were confirmed by Sanger sequencing (Fig. 1B). Indeed, all *HP6*^*CR*^ homozygous mutant flies contained two edited alleles with complete excision of the *HP6* coding region. Excision of the 3xP3-dsRED marker cassette using PBac transposase yielded viable homozygous *HP6*^*CR*^ null mutant flies indicating that the presence of the 3xP3-dsRED marker cassette did not impact viability (Gratz et al. 2015).

**Figure 1:**
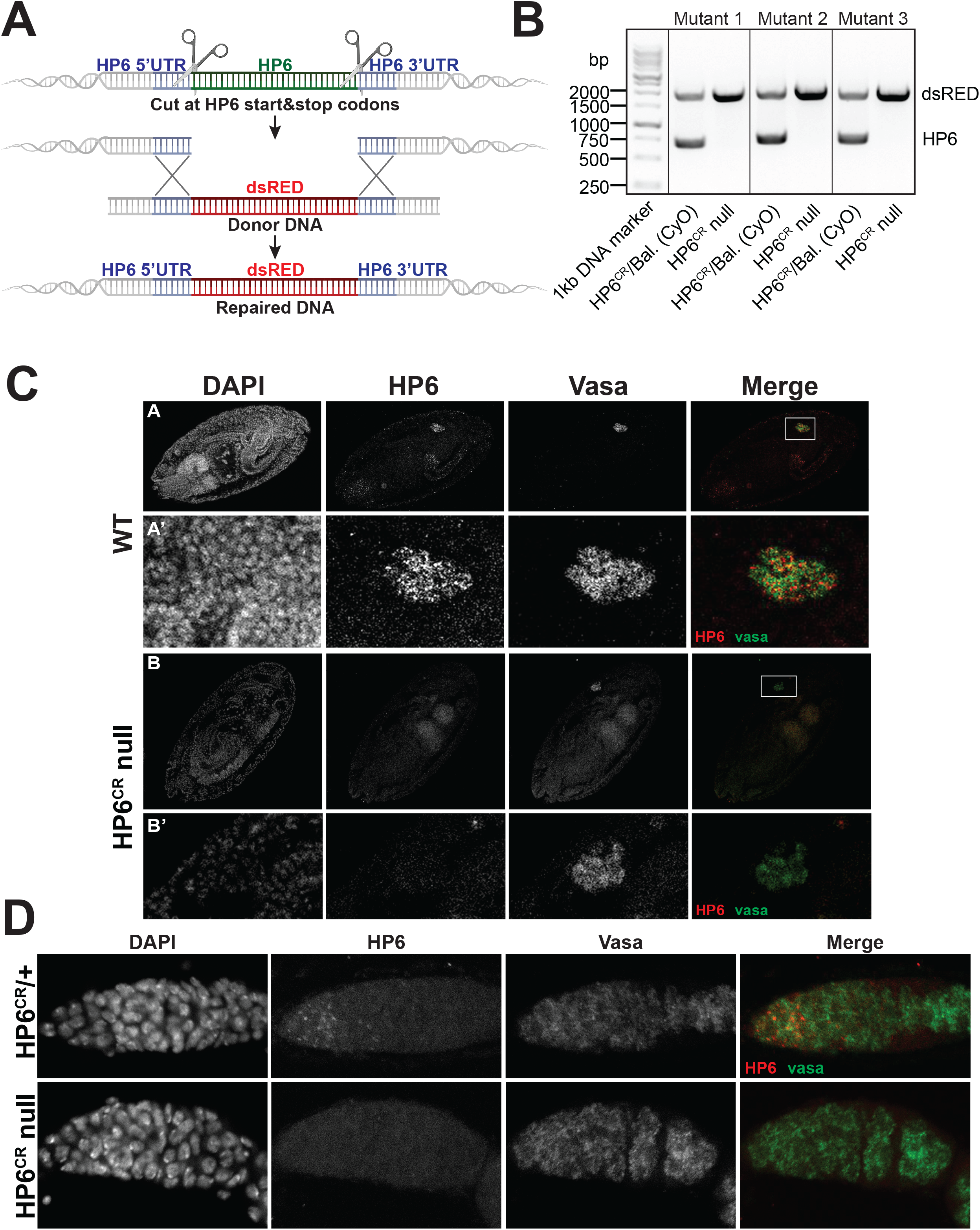
Generation of *HP6*^*CR*^ null alleles and *HP6* RNA expression in Drosophila germ cells. A) Scheme for generating an *HP6* knockout using CRISPR-mediated genome editing. Two guide RNAs were used to generate cuts near the HP6 start codon and stop codon. A repair template containing homology to the *HP6* 5’ and 3’ UTRs was used to replace the *HP6* coding region with the *3xP3-dsRED* marker cassette flanked by PBac transposon ends. B) Genomic DNA was isolated from *HP6*^*CR*^ heterozygous and homozygous mutant flies and the *HP6* genomic locus was amplified using primers flanking *HP6* in the first intron of *dumpy*. The presence of *HP6* and the *3xP3-dsRED* cassette was identified by size differences on an agarose gel and confirmed by sequencing. C) *HP6* RNA is expressed in primordial germ cells in the developing embryo. mRNA in situ hybridization using HCR in WT and *HP6*^*CR2*^ mutant embryos (as shown previously in (Li et al. 2021)). *HP6* mRNA is present in primordial germ cells (marked by Vasa) in WT stage 14 embryos but absent in *HP6*^*CR2*^ mutant embryos. White boxes encompass germ cells and indicate boundaries for the enlarged views shown in a’ and b’. A’ and B’) enlarged view of the primordial germ cells from (A) and (B) respectively. D) mRNA in situ hybridization by HCR in *HP6*^*CR2*^ heterozygous and *HP6*^*CR1*^*/HP6*^*CR2*^ trans-heterozygous mutant ovaries. *HP6* is expressed in the germline stem cells, 2-cell cysts, and 4-cell cysts controls but not in the mutants (as shown previously in (Slaidina et al. 2021)).

### *HP6* RNA is expressed in primordial germ cells and germline stem cells

To probe whether our HP6 mutants lacked HP6 transcript *in vivo*, we used in situ hybridization chain reaction (HCR) to visualize the expression of HP6 in the germline in both heterozygous and *HP6*^*CR*^ null germ cells (Fig. 1C&D). *HP6* RNA was previously identified in the PGCs of embryos and germline stem cells of adult ovaries (Jevitt et al. 2020; Rust et al. 2020; Slaidina et al. 2020; Li et al. 2021; Slaidina et al. 2021). In wildtype embryos, *HP6 i*s expressed during gonad coalescence (Li et al. 2021). Consistent with the complete deletion of the HP6 locus, we could not visualize *HP6* RNA in the PGCs of *HP6*^*CR2*^ null mutant embryos or the germline stem cells of *HP6*^*CR1*^/*HP6*^*CR2*^ trans-heterozygous mutant ovaries (Fig. 1C&D). Together, these data support previous reports of *HP6* expression in germ cells and confirm that our *HP6*^*CR*^ null mutants lack *HP6* transcripts.

### *HP6*^*CR*^ mutants are viable and fertile

Due to the significant expression of *HP6* in primordial germ cells and germline stem cells, we asked what function *HP6* has in germline development. We characterized *HP6*^*CR2*^ null mutant females for egg-laying and found that deletion of the *HP6* locus does not affect egg-laying compared to control flies (Fig. 2A). We next investigated the hatchability of eggs laid by *HP6*^*CR2*^ null mutant females. We observed that eggs laid by *HP6*^*CR2*^ mutant mothers hatched at the same frequency as controls (Fig. 2B). Our results indicate that *HP6* is not required for germline development, fertility, or viability, even when mutants were maintained in homozygosity for many generations (data not shown).

**Figure 2:**
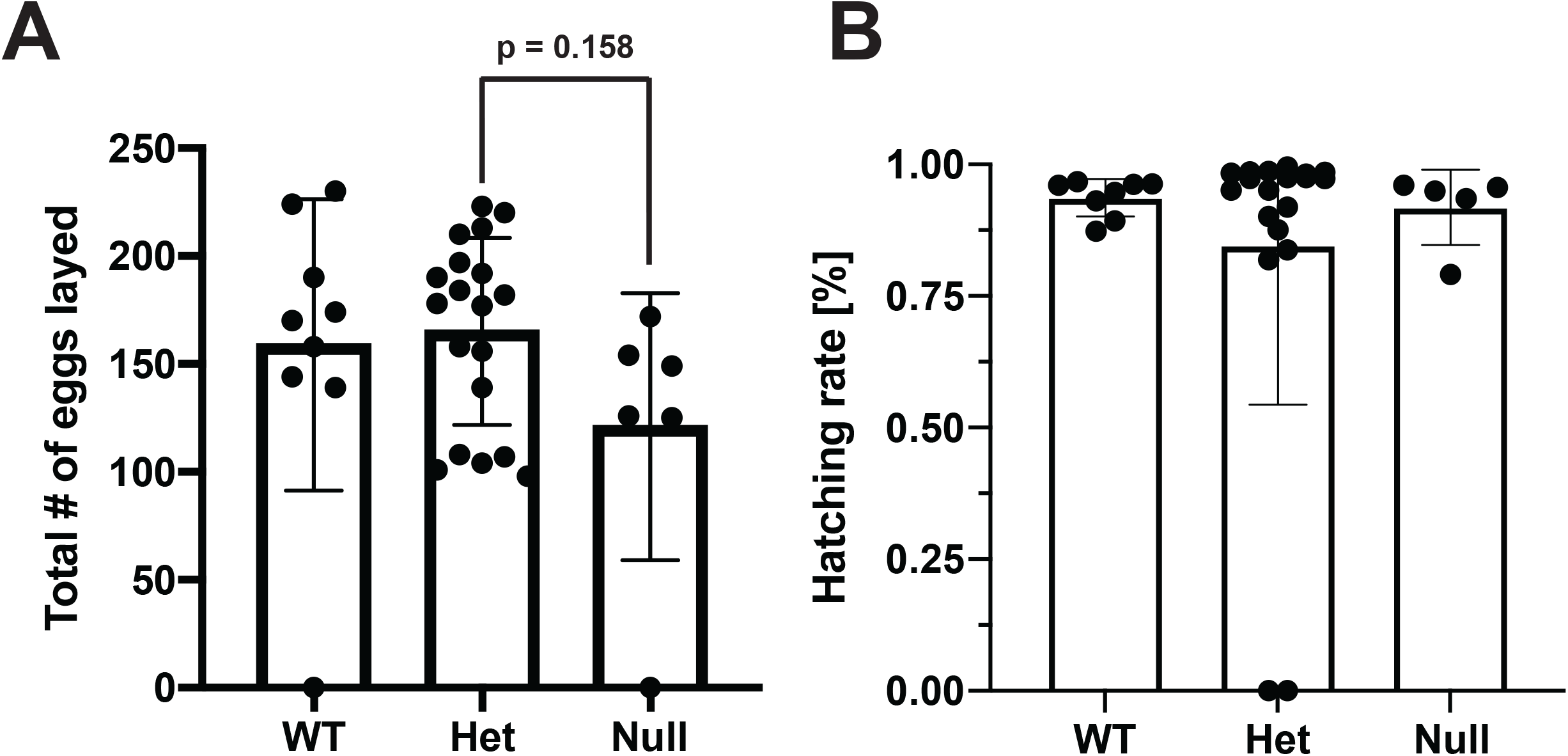
HP6 mutants are viable and fertile. A) Female fertility test for *HP6*^*CR2*^ mutants. Total number of eggs laid per female over 72 hours in wildtype (WT), *HP6*^*CR2*^ heterozygous (Het), and *HP6*^*CR2*^ null (Null) mutants. B) Hatching rate of eggs laid in (A).

### Reported *HP6* mutant chromosome harbors an unlinked, lethal mutation

Previously generated *HP6* mutants were reported to have 100% late larval-pupal lethality in *Drosophila melanogaster* (Ross et al. 2013). These mutants were reported to be generated by imprecise excision of a P-element that was inserted into the *HP6* locus. Imprecise excision of the P-element line P(GT1)HP6^[BG01429]^ reportedly yielded two mutants: HP6^366^, a complete deletion of the *HP6* locus, and HP6^35^, which encodes a mutant HP6 protein lacking a nuclear localization signal (NLS) (Ross et al. 2013). To understand the differences between our *HP6*^*CR*^ mutants and the previously generated *HP6*^*366*^ and *HP6*^*35*^ lines, we asked if our *HP6*^*CR1*^ mutant could complement the lethality of either the *HP6*^*366*^ or *HP6*^*35*^ mutants. As previously reported, *HP6*^*366*^ and *HP6*^*35*^ mutants fail to complement for lethality, indicating that the *HP6*^*366*^*/HP6*^*35*^ trans-heterozygotes are not viable (Fig. 3A). In contrast, our *HP6*^*CR1*^ null mutant was viable in trans to the *HP6*^*366*^ and *HP6*^*35*^ mutants, suggesting that the lethality of the previously generated mutants may not be caused by mutations in the *HP6* locus (Fig. 3A). In support of this conclusion, Muakai et al. generated an *HP6* mutation by P-element excision of the parental P-element line P(GT1)HP6^[BG01429]^, the same line that was used to generate *HP6*^*366*^ and *HP6*^*35*^, and found that the excision allele is viable in trans to a deletion of the locus (Mukai et al. 2007).

**Figure 3:**
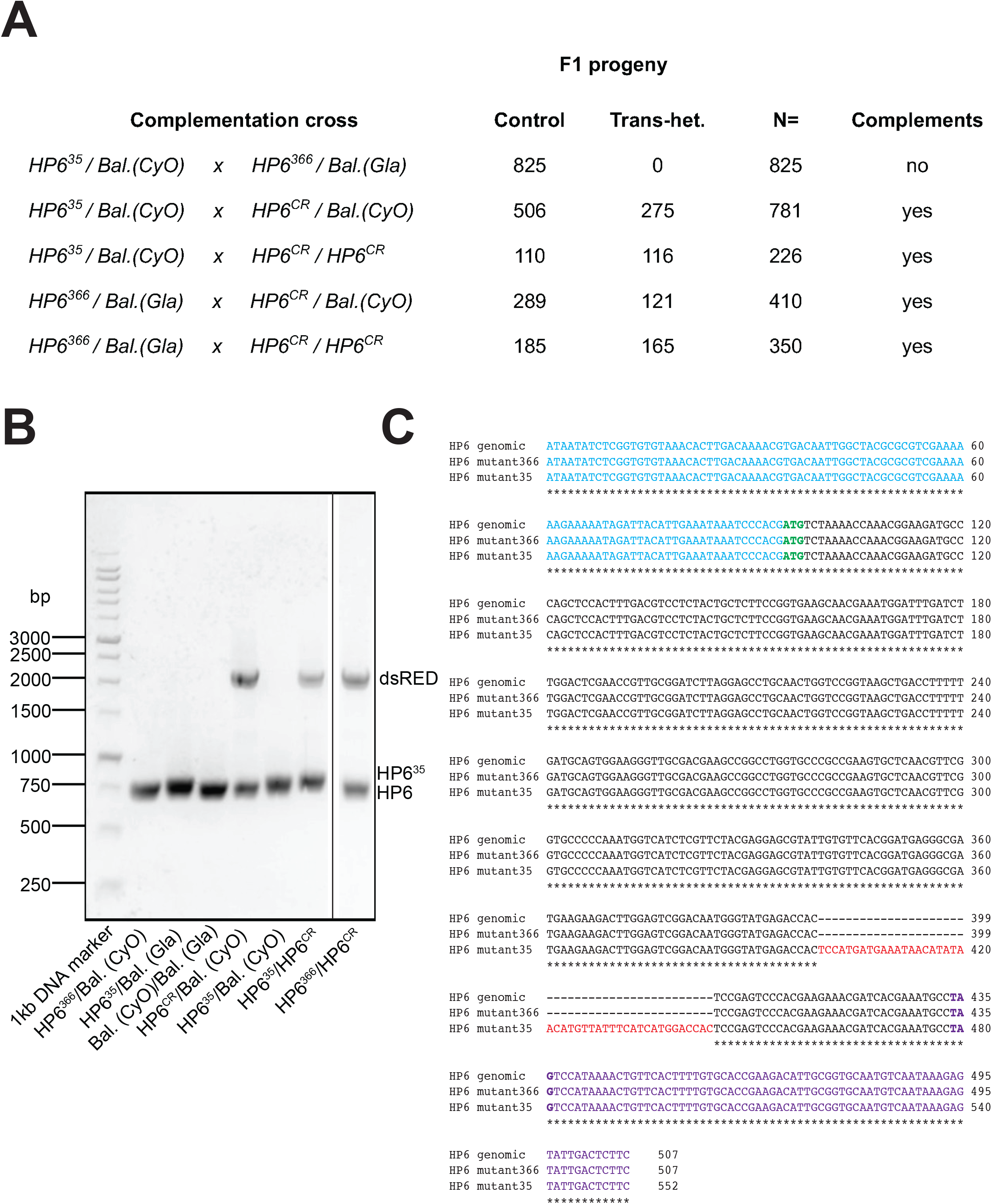
*HP6*^*35*^ and *HP6*^*366*^ lethality is independent of the *HP6* locus. A) Genetic complementation analysis for lethality between *HP6*^*CR1*^ mutants and *HP6*^*35*^ or *HP6*^*366*^ mutants. Total progeny counts are shown next to the parental genotypes for each cross. Balancer chromosomes are annotated in parental genotypes. Control progeny are described as all progeny containing a balancer chromosome. Trans-het. progeny are progeny containing two mutant HP6 chromosomes. B) Genomic DNA was isolated from the F1 progeny of crosses in A and was used to amplify the HP6 genomic locus by PCR. The presence of *HP6*^*35*^, *HP6*^*366*^, the dsRED cassette (from *HP6*^*CR1*^ mutants), and WT *HP6* from balancer chromosomes was identified by size differences on an agarose gel and confirmed by Sanger sequencing. C) Sequence alignment of *HP6*^*35*^ and *HP6*^*366*^ mutants as compared to the WT *HP6* locus. “*” beneath sequence alignments indicate identical residues. HP6^35^ contains the previously described mutation causing a premature stop codon plus additional sequence insertions in the HP6 coding region (mutation shown in red). HP6^366^ contains a WT copy of HP6 rather than the previously described complete deletion of the *HP6* locus. The HP6 5’UTR is shown in light blue, the start codon is shown in green, the stop codon is shown in bolded purple, and the 3’UTR is shown in purple.

To confirm that crosses between our *HP6*^*CR*^ alleles and the *HP6*^*366*^ or *HP6*^*35*^ mutants resulted in viable trans-heterozygous progeny containing two HP6 mutant alleles, we isolated genomic DNA from individual progeny and amplified the *HP6* locus by PCR. The amplified products were run on an agarose gel and alleles were identified by differences in size prior to sequencing (Fig. 3B&C). All balancer chromosomes contained a wildtype copy of the *HP6* locus, while the *HP6*^*CR*^ alleles contained the inserted 3xP3-dsRED marker cassette in place of the *HP6* coding region (Fig. 3B). As had been reported previously, the *HP6*^*35*^ allele contained a 45 base pair insertion that resulted in a premature stop codon at the 97^th^ amino acid of the HP6 protein, prior to the predicted nuclear localization signal (Fig. 3B&C) (Ross et al. 2013). Analysis of the *HP6*^*366*^ mutant, however, revealed a wildtype sequence for the *HP6* locus, rather than the reported complete deletion of the *HP6* gene (Fig. 3B&C) (Ross et al. 2013). Despite this, the chromosome bearing the *HP6*^*366*^ mutant is homozygous lethal and lethal in trans to the chromosome bearing the *HP6*^*35*^ allele (Fig. 3A). We concluded that the lethality associated with the *HP6*^*366*^ and *HP6*^*35*^ alleles is independent of the *HP6* locus.

### Unlinked lethal mutation is not associated with the *HP6* locus

To further probe whether the lethality of the *HP6*^*35*^ and *HP6*^*366*^ mutants is a result of a loss of functional *HP6*, we sought to rescue *HP6* expression of the *HP6*^*35*^*/HP6*^*366*^ trans-heterozygotes by expressing an HP6 wildtype transgene. Since the pUASp:HP6-GFP transgene generated by Ross et al. was inserted on the second chromosome at an unknown location, we generated recombinants between the *HP6*^*35*^ mutant line and the *pUASp:HP6-GFP* transgene (Ross et al. 2013). The crossing scheme for the generation of recombinants between the *HP6*^*35*^ mutant and the *pUASp:HP6-GFP* transgene is shown in Figure 4A. We generated recombinant lines that were scored for the *pUASp:HP6-GFP* transgene, which carries the marker *w+* and produces red eye color. Once recombinant lines were isolated, we used PCR of the *HP6* locus to detect the *HP6*^*35*^ mutation. Subsequently, all recombinant lines were crossed with the *HP6*^*366*^ mutant line to generate *UAS:HP6-GFP, HP6*^*35*^*/HP6*^*366*^ trans-heterozygotes (Fig. 4A). A total of eighty-eight potential recombinant lines were isolated based on the presence of red eyes due the *UAS:HP6-GFP* transgene. Of these, sixty had the parental genotype, containing the *UAS:HP6-GFP* transgene and a wildtype *HP6* locus that resulted in fully viable trans-heterozygotes in trans to *HP6*^*366*^. Eighteen recombinant lines were lethal in trans to the *HP6*^*366*^ mutant (Fig. 4B, Supplemental File 1); of these, four had a wildtype *HP6* locus. Twenty-four lines contained the *HP6*^*35*^ mutation, of these, ten were viable in trans to the *HP6*^*366*^ allele (Fig. 4B). Finally, all recombinants were also kept as balanced stocks, creating the opportunity to produce homozygous mutant progeny. All recombinants that were viable in trans to the *HP6*^*366*^ allele were also viable as homozygotes. In contrast, recombinants that were lethal in trans to the *HP6*^*366*^ allele were also homozygous lethal in the stock, irrespective of the presence of the *HP6*^*35*^ mutation (Supplemental File 1). We conclude that the lethality associated with the *HP6*^*35*^ and *HP6*^*366*^ alleles was due to an unlinked lethal mutation that maps between the *UAS:HP6-GFP* transgene and the *HP6* locus.

**Figure 4:**
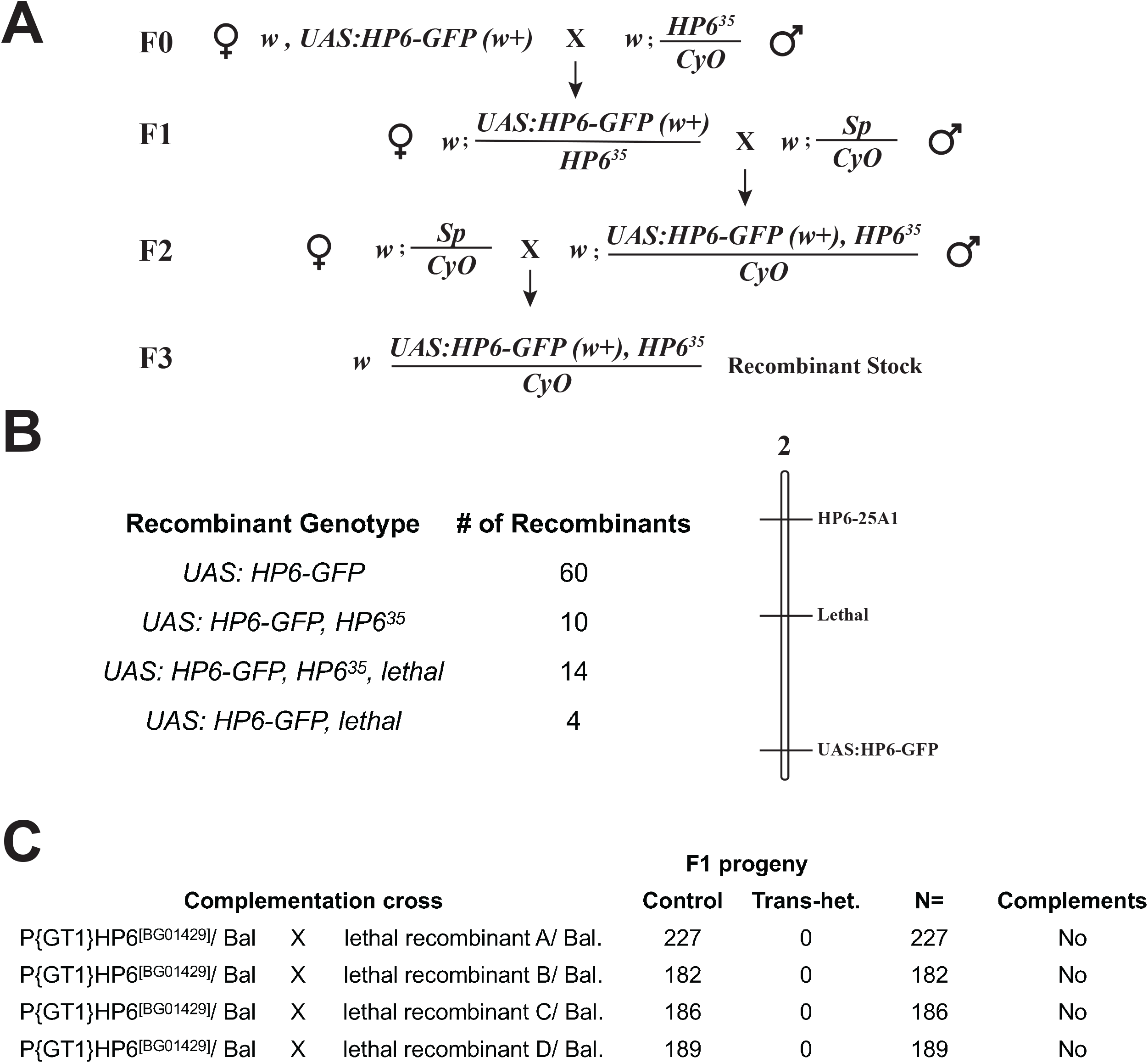
Identification of an independent lethal mutation in the *P(GT1)HP6*^*[BG01429*]^ stock. A) Genetic crossing scheme for the generation of recombinants between *HP6-GFP* and *HP6*^*35*^. Female genotypes are shown on the left, male genotypes are shown on the right. B) Genetic mapping of an unlinked lethal mutation. Recombinants were scored for: *HP6-GFP* (by red eye color), the *HP6*^*35*^ mutation (by PCR genotyping), and the lethal mutation (by complementation analysis for lethality with the *HP6*^*366*^ chromosome). Linkage map is shown on the right. For a list of recombinants see Supplemental File 1. C) Genetic complementation analysis for lethality with *P(GT1)HP6*^*[BG01429*]^. None of the four recombinant lines that contain the lethal mutation but a WT HP6 locus complemented the *P(GT1)HP6*^*[BG01429*]^ line, indicating that the lethal mutation is present in the parental *P(GT1)HP6*^*[BG01429*]^ line.

Our data were also consistent with the notion that this lethal mutation was already present on the parental P(GT1)HP6^[BG01429]^ chromosome prior to the generation of the *HP6*^*35*^ and *HP6*^*366*^ alleles. To test this idea, we performed complementation analysis for lethality between the parental P(GT1)HP6^[BG01429]^ line and our four recombinant lines that had a wildtype *HP6* locus but contained the lethal mutation (Fig. 4C). None of the four recombinants were viable in trans to the parental P-element line. This suggests that the unlinked mutation was not acquired during the P-element excision process, but instead was already present in the parental line when the *HP6*^*35*^ and *HP6*^*366*^ mutants were generated.

Our genetic analysis demonstrates that a lethal mutation was present on the parental chromosome of the *P(GT1)HP6*^*[BG01429*]^ line prior to the generation of the *HP6*^*35*^ and *HP6*^*366*^ mutants, preventing accurate interpretation of the essentiality of the *HP6* gene. The problems highlighted here could have been avoided by genetic analysis of P-element excision mutants, such as the use of precise P-element excisions to ascertain that the lethality of insertion is caused by the P-element in question. Indeed, the *HP6*^*366*^ mutant, which has a wildtype *HP6* locus, may have been the result of precise excision of the P-element. Alternatively, when a precise P-element excision allele is unavailable, P-elements can be tested in trans to an independent line, such as a deletion for the locus as was done in Mukai et al. (Mukai et al. 2007).

### RNAi-mediated lethality is HP6 independent

In addition to genetic mutants, previous studies also used RNAi-based methods to probe *HP6* function *in vivo* (Joppich et al. 2009; Chen et al. 2010; Ross et al. 2013). Each of these studies found that RNAi-mediated knockdown of *HP6* resulted in lethality. To ask if the RNAi-mediated lethality was due to the loss of HP6 or to an off-target effect, we expressed *HP6* RNAi in *HP6*^*CR1*^ heterozygous and null mutant flies (Fig 5A&B). Since our *HP6*^*CR1*^ mutant lines are deleted for the complete *HP6* coding sequence, they should lack the sequences targeted by the RNAi transgene. In previous studies three GD lines were generated by P-element insertion, each containing the same inverted repeat DNA fragment that is homologous to the coding region of the *HP6* gene (Dietzl et al. 2007). This transgene was inserted into three separate genomic locations, generating three RNAi lines: GD13072, GD13073, and GD13074. As in previous studies, we drove the RNAi transgene with a ubiquitous GAL4 driver, *tubulin*:GAL4. Consistent with prior reports, GD13072 and GD13073 expression caused lethality in *HP6*^*CR*^ heterozygous mutant flies (Fig. 5A&B) (GD13074 is no longer available). Notably, the RNAi also resulted in lethality in *HP6*^*CR1*^ null mutants, despite them lacking all *HP6* locus homology regions. Together these data point to an off-target, lethal effect of the RNAi. As a control for RNAi specificity, Ross et al. had shown partial rescue of the RNAi-induced lethality by expressing a *UAS:HP6-GFP* transgene. While it is possible that the lethality of the RNAi expression may have been titrated by *HP6* wildtype sequences expressed at higher levels from the transgene than the endogenous locus, it is more likely that this partial rescue was caused by reduced expression of the RNAi transgene due to splitting GAL4-driver activity between two UAS constructs (*UAS:HP6-GFP* and *UAS-RNAi*). Together our data demonstrate that the RNAi transgene results were due to a lethal off-target effect that is independent of the *HP6* locus.

**Figure 5:**
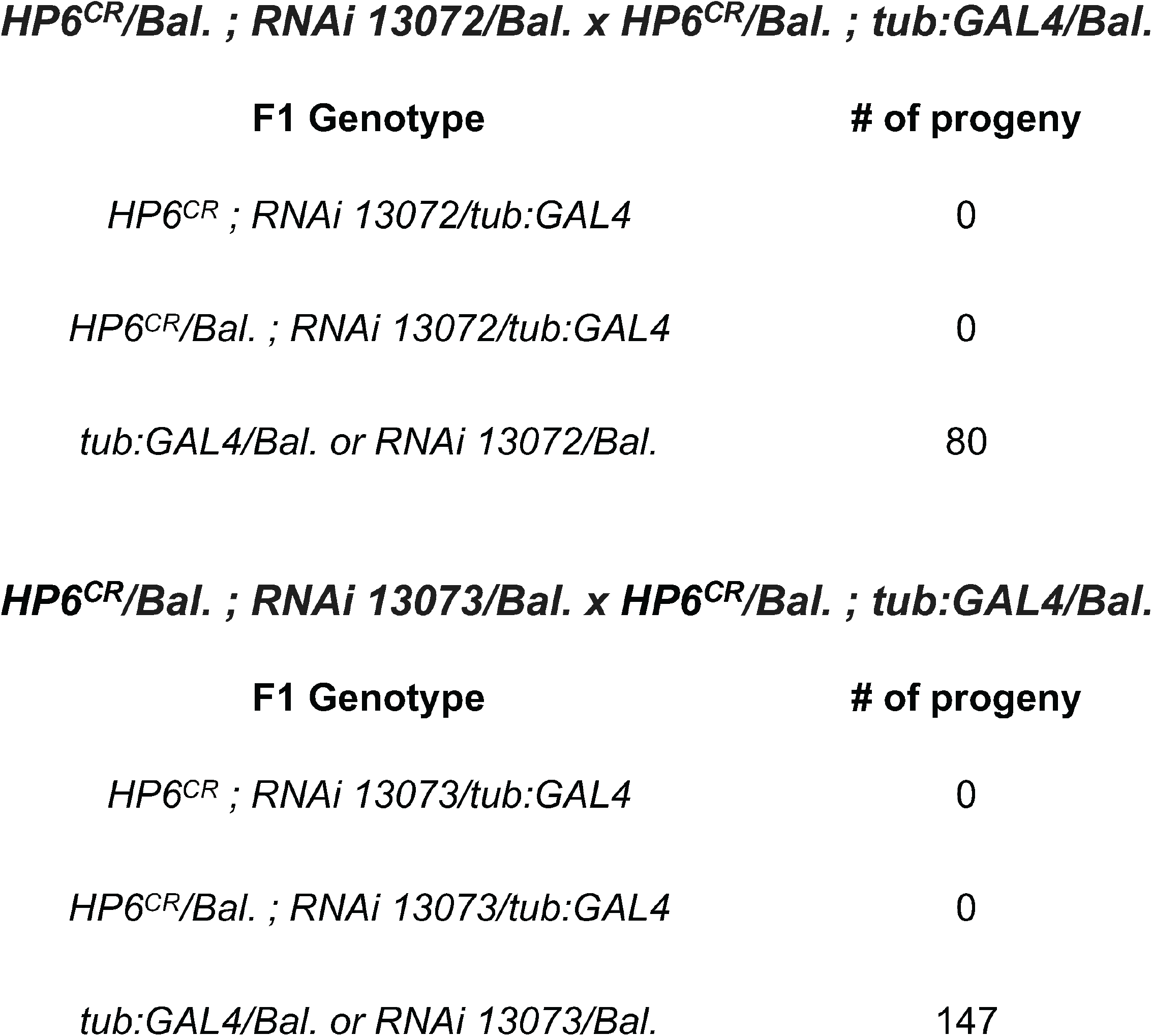
RNAi-mediated lethality is *HP6* independent. Interfering RNA-13072 (top) and RNA-13073 (bottom) complementary to *HP6* cause lethality when driven with a ubiquitous *tubulin* promoter (*tubulin*:GAL4). *HP6*^*CR*^ null mutants are sensitive to RNAi despite lacking *HP6* mRNA, indicating an off-target, lethal effect of the RNAi.

While screens with available reagents such as RNAi have great potential to identify candidate genes, thorough genetic analysis is necessary to validate these findings. When using RNAi, it is advisable to use several independent RNAi insertions, and controls should be employed to account for possible titration effects of the Gal4 activity when driving multiple UAS constructs. While more precise, CRISPR-based mutants may also be insufficient to conclusively probe phenotype, because of the nature of their genetic defects. Aberrant mRNAs generated from mutagenesis can trigger a compensatory increase in the expression of redundant genes with high sequence similarity, termed transcriptional adaptation (Jakutis and Stainier 2021). However, this compensatory mechanism cannot explain the viability of the *HP6*^*CR*^ mutants. First, *HP6* has minimal sequence similarity to any other related gene, including *HP1B*, that is necessary for compensation (Levine et al. 2012; Ross et al. 2013) and second, *HP6*^*CR*^ mutants do not encode any degradation defective RNA that is necessary to trigger this adaptation (El-Brolosy et al. 2019; Jakutis and Stainier 2021).

Our data call into question the idea that up to 30% of young genes can quickly acquire essential functions (Chen et al. 2010; Xia et al. 2021). Additional studies have recently characterized the essentiality of young genes by performing in-depth functional studies using newer CRISPR-based mutagenesis methods (Kondo et al. 2017; Lee et al. 2019; Kasinathan et al. 2020). We compiled the current knowledge of the 61 young essential genes that were identified in a previous RNAi screen (Chen et al. 2010) (Supplemental File 2). Many of these genes were tested for essentiality with a single RNAi line and the majority of those deemed essential were tested with RNAi constructs inserted at chromosome position 40D, an insertion site that results in lethality when ubiquitously expressed irrespective of the RNAi used (Green et al. 2014). Since then, additional studies have found that the vast majority of the originally identified young essential genes are not essential when tested with orthologous methods (Supplemental File 2). Taken together, these findings suggest that the percentage of young genes that gain essential function is ten times lower than previously reported, and is closer to 3 percent rather than the predicted 30 percent. Importantly, a few cases of young essential genes have been validated (Chen et al. 2010; Kasinathan et al. 2020), confirming that young genes can gain essential function, though they are likely the exception rather than the rule. Together, our data highlight how multiple orthogonal approaches are essential to uncover the true proportion of young genes that have gained an essential function.

## Materials and Methods Nomenclature

Throughout the text, we use *HP6* to refer to the gene which is also known as *Umbrea* and *CG15636* to follow FlyBase guidelines.

### Fly stocks and husbandry

All stocks and crosses were maintained at 18°C and 25°C respectively, on cornmeal molasses yeast medium. *Umbrea*^*35*^*(HP6*^*35*^*), Umbrea*^*366*^ (*HP6*^366^) and *Umbrea-GFP* (*pUASp:HP6-GFP*) lines were gifts from Harmit Malik (Fred Hutchinson Cancer Research Center). The *HP6*^*CR*^ lines were generated for this study as described below. Lehmann lab Oregon-R (OrR) stocks were used in fertility assays and hatching rate experiments. *HP6* RNAi lines 13072 and 13073 were purchased from the Vienna Drosophila Resource Center (VRDC) and were used in RNAi experiments alongside a Lehmann lab *tubulin*:GAL4 driver line.

### Generation of *HP6* mutant alleles

For the generation of *HP6*^*CR1*^, CRISPR guides AGGACGTCAAAGTGGAGCT and ATCGTTTCTTCCTGGGACT were cloned into the pU6 tandem gRNA vector (DGRC #1436) and injected into w[1118]; PBac{y[+mDint2]=vas-Cas9}VK00027/TM6B, Tb[1] embryos carrying the *vas-Cas9* transgene by Bestgene. The pHD-ScarlessDsRed donor vector (DGRC #1364) was used as a repair template. A 920 base pair upstream homology arm was cloned upstream of the 3xP3-dsRED marker cassette beginning at -860 upstream of the HP6 +1 site and ending at +56 within the HP6 5’UTR. A 1004 base pair downstream homology arm was cloned downstream of the 3xP3-dsRED marker cassette beginning at the *HP6* stop codon. Use of this repair template resulted in excision of the *HP6* coding region and insertion of the 3xP3-dsRED marker cassette flanked by PBac transposon ends. A single, edited founder male was isolated, the Cas9 was removed from the genome, and an isogenic balanced stock was generated. For the generation of HP6^CR2^ mutant line, CRISPR guides AGGACGTCAAAGTGGAGCT and AAGACTTGGAGTCGGACAA were cloned in to the pU6 tandem gRNA vector (DGRC #1436) and injected into y[1] sc[*] v[1] sev[21];{nos-Cas9}attP2 embryos carrying the *nos-Cas9* transgene by Bestgene. The same donor vector was used as a repair template. The Cas9 was removed from the genomes, and isogenic balanced stocks were generated. Six additional CRISPR-generated mutant lines were also isolated but were not used in this study.

### Genomic DNA extraction and genotyping of HP6 mutants

Genomic DNA was extracted from individual adult *D. melanogaster* flies. Single flies were homogenized in 50 uL of buffer containing 10 mM Tris pH 8, 1 mM EDTA and 25mM NaCl. To each homogenate, Proteinase K was added to a final concentration of 0.2 mg/mL and samples were incubated at 55°C for 30 minutes with mixing at 800 rpm. To inactivate the Proteinase K, samples were incubated at 95°C for 3 minutes and subsequently centrifuged at 1000xg for 5 minutes. The supernatant was collected as clean genomic DNA. PCR reactions were performed using primers flanking the HP6 locus in the first intron of the *dumpy* gene. For genotyping of HP6^CR^ mutants, primers HSM208: CAACCAGTTCGCATGAAAATGCATAATCAATC and HSM215: AGATGAAGATGCACCAATGATACGCTCAATGGG were used (Ross et al. 2013). PCR products were run on an agarose gel, each product was isolated, purified, and Sanger sequenced to confirm the presence/absence of a wildtype or mutant *HP6* locus.

### RNA in situ hybridization

For ovary HCR, the protocol was adapted from (Choi et al. 2018) and performed exactly as previously described (Slaidina et al. 2021). The HCR protocol for use in Drosophila embryos was adapted from (Trcek et al. 2017) and Molecular Instruments, Inc. Briefly, embryos were collected, fixed, permeabilized, and postfixed exactly as in (Trcek et al. 2017). Embryos were washed with PBS-0.1% Tween, pre-hybridized for 30 minutes at 37°C and incubated overnight with 0.8-1.6 pmol of custom *HP6* probe (Molecular Instruments, Inc) at 37°C. Embryos were then washed and probes were amplified using appropriate hairpins that had been heat shocked at 95°C for 90 seconds and allowed to slow cool at room temperature for 30-90 minutes. Hairpins were incubated with embryos overnight at room temperature before excess hairpins were washed off using 5x SSC containing 0.1% Tween. Embryos were equilibrated in Vectashield overnight at 4°C and imaged using standard confocal fluorescence microscopy. Custom-designed probes, probe hybridization buffer, probe wash buffer, and amplification buffer were procured from Molecular Instruments, Inc.

### Fertility and viability assays

Fertility and viability assays were performed with *D. melanogaster HP6*^*CR*^ null mutant flies. To obtain control heterozygous siblings, we generated heterozygous *HP6*^*CR*^ mutants without the CyO balancer, as this balancer results in a slight decrease in fertility (data not shown). To obtain *HP6*^*CR*^ homozygous mutant and control heterozygous siblings, the following crosses were performed: *HP6*^*CR*^/CyO heterozygous virgin females were crossed with OrR males. F1 straight-wing males and virgin females were crossed to generate F2 virgin females that were used in fertility assays. F2 virgin females were collected for two days. Females were subsequently aged for 3 days at 25°C. To test for fertility, crosses were performed with 1 female and 2 OrR males. Crosses were flipped to a fresh apple juice plates with fresh yeast every 24 hours and embryos were counted. Apple juice plates remained at 25°C for an additional 24 hours and were counted a second time for the presence of hatched and unhatched embryos. After completion of the 72 hours, females were collected and genotyped using PCR for the presence of *HP6* and dsRED to identify *HP6*^*CR*^ null mutants, heterozygous, and wildtype controls. Plots were made in Prism and student t-tests were performed to generate p-values. The total number of eggs laid was used as a measure of fertility and hatch rate was calculated as the total number hatched embryos divided by the total number of laid embryos per day.

### Generation of *HP6* recombinants

Females carrying the *pUASp-HP6-GFP* transgene, which contains a *w+* gene and is referred to as *HP6-GFP* females, were crossed with *HP6*^*35*^ heterozygous males to produce *HP6-GFP/HP6*^*35*^ F1 progeny (see crossing scheme Figure 4A). F1 *HP6-GFP/HP6*^*35*^ virgin females were collected and crossed with males containing a second chromosome dominant marker (Sp) in trans to the CyO balancer chromosome. All F2 red-eyed males were collected and individually crossed with Sp/CyO virgin females to generate CyO-balanced isogenic, recombinant lines in the F3. F2 crosses were allowed to lay until larvae were visible in vials and then the F2 males were removed and subsequently used for genomic DNA isolation and genotyping of the *HP6* locus by PCR. For genotyping recombinants, primers HSM215:AGATGAAGATGCACCAATGATACGCTCAATGGG and HP6_96-104Rev: CGTGATCGTTTCTTCGTGGGAC were used. PCR products were subjected to gel electrophoresis and the presence or absence of the *HP6*^*35*^ mutation was determined by size. Males from the F3 recombinant lines were then crossed with *HP6*^*366*^ virgin females to assess HP6^366^ complementation for lethality.

### *HP6* RNAi

*HP6* RNAi lines 13072 and 13073 and the Tub-GAL4 chromosomes were crossed into *HP6*^*CR*^ mutant background. F1 progeny were isolated and counted for the presence of homozygous or heterozygous *HP6*^*CR*^ null mutants containing both the RNAi and Tub-GAL4 chromosomes. F1 Progeny containing the RNAi or the Tub-GAL4 chromosome, but not both, were used as controls. Crosses were maintained at 25°C throughout.

## Author Contributions

S.G. and R.L. conceived the experiments and wrote the manuscript with helpful feedback from A.R. S.G. conducted all experiments with help from A.R. and M.S. R.L. and S.G. acquired funding for the project.

## Acknowledgements

We want to thank Dr. Harmit Malik (Fred Hutchinson Cancer Center) for his advice, support, and encouragement, and the Malik lab for sharing reagents. We want to thank the Lehmann lab, particularly Arjuna Rajakumar and Julia Wucherpfennig, for their helpful feedback, discussions, and comments on the manuscript. Transgenic fly stocks were obtained from the Vienna *Drosophila* Resource Center (VDRC, http://www.vdrc.at). S.G. is supported by the American Cancer Society Postdoctoral Fellowship PF-21-116-01-RMC. R.L. is supported by R37 HD41900.

## Supplemental File legends

Supplemental File 1: Genotype, viability, and results of the complementation analysis for lethality with *HP6*^*366*^ for each *HP6* recombinant line. Recombinant stocks highlighted in red contain both the lethal mutation and the *HP6*^*35*^ mutation and fail to complement lethality associated with *HP6*^*366*^ chromosome. Recombinant lines that are highlighted in green contain the *HP6*^*35*^ mutation, are homozygous viable, and viable in trans to HP6^366^ chromosome. Stocks highlighted in blue contain the lethal mutation but not the *HP6*^*35*^ mutation and are lethal in trans to HP6^366^ chromosome.

Supplemental File 2: Significantly fewer young genes gain essential function than previously thought. Analysis of 59 young genes identified in Chen et al. as essential through RNAi knockdown with the VRDC KK library (Chen et al. 2010). The original VRDC KK RNAi line that was used for each knockdown in the Chen et al. study is shown by VRDC ID. Presence/absence of the 40D insertion site that results in lethality when ubiquitously expressed is noted for each RNAi line that was previously used (Dietzl et al. 2007; Green et al. 2014). Our current understanding of the essentiality status for each gene based on available literature is noted. Genes were labeled as viable if an orthologous method, such as CRISPR mutagenesis or knockdown with a second, independent RNAi construct, resulted in viable animals. Genes were labeled as lethal (highlighted in red) if two or more orthologous methods, such as RNAi and CRISPR mutagenesis, both resulted in lethality or if two or more distinct RNAi lines resulted in lethality. Genes highlighted in yellow have not been tested with an additional RNAi construct or orthologous method, to our knowledge. Of the 59 young genes that were identified as essential, 54 are viable when tested with additional methods, two are lethal.

## References

Chen S, Krinsky BH, Long M. 2013. New genes as drivers of phenotypic evolution. Nat Rev Genet 14: 645–660.

Chen S, Zhang YE, Long M. 2010. New genes in Drosophila quickly become essential. Science 330: 1682–1685.

Choi HMT, Schwarzkopf M, Fornace ME, Acharya A, Artavanis G, Stegmaier J, Cunha A, Pierce NA. 2018. Third-generation. Development 145.

Dietzl G, Chen D, Schnorrer F, Su KC, Barinova Y, Fellner M, Gasser B, Kinsey K, Oppel S, Scheiblauer S et al. 2007. A genome-wide transgenic RNAi library for conditional gene inactivation in Drosophila. Nature 448: 151–156.

Ding Y, Zhao L, Yang S, Jiang Y, Chen Y, Zhao R, Zhang Y, Zhang G, Dong Y, Yu H et al. 2010. A young Drosophila duplicate gene plays essential roles in spermatogenesis by regulating several Y-linked male fertility genes. PLoS Genet 6: e1001255.

El-Brolosy MA, Kontarakis Z, Rossi A, Kuenne C, Günther S, Fukuda N, Kikhi K, Boezio GLM, Takacs CM, Lai SL et al. 2019. Genetic compensation triggered by mutant mRNA degradation. Nature 568: 193–197.

Gratz SJ, Rubinstein CD, Harrison MM, Wildonger J, O’Connor-Giles KM. 2015. CRISPR-Cas9 Genome Editing in Drosophila. Current protocols in molecular biology / edited by Frederick M Ausubel [et al] 111: 31.32.31-31.32.20.

Green EW, Fedele G, Giorgini F, Kyriacou CP. 2014. A Drosophila RNAi collection is subject to dominant phenotypic effects. Nat Methods 11: 222–223.

Greil F, de Wit E, Bussemaker HJ, van Steensel B. 2007. HP1 controls genomic targeting of four novel heterochromatin proteins in Drosophila. EMBO J 26: 741–751.

Haerty W, Jagadeeshan S, Kulathinal RJ, Wong A, Ravi Ram K, Sirot LK, Levesque L, Artieri CG, Wolfner MF, Civetta A et al. 2007. Evolution in the fast lane: rapidly evolving sex-related genes in Drosophila. Genetics 177: 1321–1335.

Ida H, Suzusho N, Suyari O, Yoshida H, Ohno K, Hirose F, Itoh M, Yamaguchi M. 2009. Genetic screening for modifiers of the DREF pathway in Drosophila melanogaster: identification and characterization of HP6 as a novel target of DREF. Nucleic Acids Res 37: 1423–1437.

Jakutis G, Stainier DYR. 2021. Genotype-Phenotype Relationships in the Context of Transcriptional Adaptation and Genetic Robustness. Annu Rev Genet 55: 71–91.

Jevitt A, Chatterjee D, Xie G, Wang XF, Otwell T, Huang YC, Deng WM. 2020. A single-cell atlas of adult Drosophila ovary identifies transcriptional programs and somatic cell lineage regulating oogenesis. PLoS Biol 18: e3000538.

Joppich C, Scholz S, Korge G, Schwendemann A. 2009. Umbrea, a chromo shadow domain protein in Drosophila melanogaster heterochromatin, interacts with Hip, HP1 and HOAP. Chromosome Res 17: 19–36.

Kasinathan B, Colmenares SU, McConnell H, Young JM, Karpen GH, Malik HS. 2020. Innovation of heterochromatin functions drives rapid evolution of essential ZAD-ZNF genes in. Elife 9.

Kondo S, Vedanayagam J, Mohammed J, Eizadshenass S, Kan L, Pang N, Aradhya R, Siepel A, Steinhauer J, Lai EC. 2017. New genes often acquire male-specific functions but rarely become essential in. Genes Dev 31: 1841–1846.

Lee YCG, Ventura IM, Rice GR, Chen DY, Colmenares SU, Long M. 2019. Rapid Evolution of Gained Essential Developmental Functions of a Young Gene via Interactions with Other Essential Genes. Mol Biol Evol 36: 2212–2226.

Levine MT, McCoy C, Vermaak D, Lee YC, Hiatt MA, Matsen FA, Malik HS. 2012. Phylogenomic analysis reveals dynamic evolutionary history of the Drosophila heterochromatin protein 1 (HP1) gene family. PLoS Genet 8: e1002729.

Li YR, Lai HW, Huang HH, Chen HC, Fugmann SD, Yang SY. 2021. Trajectory mapping of the early. Genome Res 31: 1011–1023.

Long M, Betrán E, Thornton K, Wang W. 2003. The origin of new genes: glimpses from the young and old. Nat Rev Genet 4: 865–875.

Loppin B, Lepetit D, Dorus S, Couble P, Karr TL. 2005. Origin and neofunctionalization of a Drosophila paternal effect gene essential for zygote viability. Curr Biol 15: 87–93.

McLysaght A, Hurst LD. 2016. Open questions in the study of de novo genes: what, how and why. Nat Rev Genet 17: 567–578.

Mukai M, Hayashi Y, Kitadate Y, Shigenobu S, Arita K, Kobayashi S. 2007. MAMO, a maternal BTB/POZ-Zn-finger protein enriched in germline progenitors is required for the production of functional eggs in Drosophila. Mech Dev 124: 570–583.

Ross BD, Rosin L, Thomae AW, Hiatt MA, Vermaak D, de la Cruz AF, Imhof A, Mellone BG, Malik HS. 2013. Stepwise evolution of essential centromere function in a Drosophila neogene. Science 340: 1211–1214.

Rust K, Byrnes LE, Yu KS, Park JS, Sneddon JB, Tward AD, Nystul TG. 2020. A single-cell atlas and lineage analysis of the adult Drosophila ovary. Nat Commun 11: 5628.

Slaidina M, Banisch TU, Gupta S, Lehmann R. 2020. A single-cell atlas of the developing. Genes Dev 34: 239–249.

Slaidina M, Gupta S, Banisch TU, Lehmann R. 2021. A single-cell atlas reveals unanticipated cell type complexity in. Genome Res 31: 1938–1951.

Trcek T, Lionnet T, Shroff H, Lehmann R. 2017. mRNA quantification using single-molecule FISH in Drosophila embryos. Nature protocols 12: 1326–1348.

VanKuren NW, Long M. 2018. Gene duplicates resolving sexual conflict rapidly evolved essential gametogenesis functions. Nat Ecol Evol 2: 705–712.

Vermaak D, Malik HS. 2009. Multiple roles for heterochromatin protein 1 genes in Drosophila. Annu Rev Genet 43: 467–492.

Witt E, Benjamin S, Svetec N, Zhao L. 2019. Testis single-cell RNA-seq reveals the dynamics of de novo gene transcription and germline mutational bias in. Elife 8.

Xia S, VanKuren NW, Chen C, Zhang L, Kemkemer C, Shao Y, Jia H, Lee U, Advani AS, Gschwend A et al. 2021. Genomic analyses of new genes and their phenotypic effects reveal rapid evolution of essential functions in Drosophila development. PLoS Genet 17: e1009654.

Yeh SD, Do T, Chan C, Cordova A, Carranza F, Yamamoto EA, Abbassi M, Gandasetiawan KA, Librado P, Damia E et al. 2012. Functional evidence that a recently evolved Drosophila sperm-specific gene boosts sperm competition. Proc Natl Acad Sci U S A 109: 2043–2048.

